# The balance N1 and the ERN correlate in amplitude across individuals in small samples of younger and older adults

**DOI:** 10.1101/2022.09.16.508295

**Authors:** Aiden M. Payne, Lena H. Ting, Greg Hajcak

**Affiliations:** Florida State University; Emory University and Georgia Tech

## Abstract

The error-related negativity (ERN) is a neural correlate of error monitoring often used to investigate individual differences in developmental, mental health, and adaptive contexts. However, limited experimental control over errors presents several confounds to its measurement. An experimentally controlled disturbance to standing balance evokes the balance N1, which we previously suggested may share underlying mechanisms with the ERN based on a number of shared features and factors. We now measure whether the balance N1 and ERN are correlated across individuals within two small groups (N=21 young adults and N=20 older adults). ERNs were measured in arrow flanker tasks using hand and foot response modalities (ERN-hand and ERN-foot). The balance N1 was evoked by sudden slip-like movements of the floor while standing. The ERNs and the balance N1 showed good and excellent internal consistency, respectively, and were correlated in amplitude in both groups. One principal component strongly loaded on all three evoked potentials, suggesting that the majority of individual differences are shared across the three ERPs. However, there remains a significant component of variance shared between the ERN-hand and ERN-foot beyond what they share with the balance N1. It is unclear whether this component of variance is specific to the arrow flanker task, or something fundamentally related to error processing that is not evoked by a sudden balance disturbance. If the balance N1 were to reflect error processing mechanisms indexed by the ERN, balance paradigms offer several advantages in terms of experimental control over errors.

## 1. Introduction

The error-related negativity (ERN) is a neural correlate of error monitoring often used to investigate individual differences in developmental, mental health, and adaptive contexts. The ERN is a negative deflection in frontocentral midline EEG after spontaneous mistakes in a variety of speeded forced-choice response tasks (Meyer, Riesel, & Proudfit, 2013; Riesel, Weinberg, Endrass, Meyer, & Hajcak, 2013). The ERN is thought to arise from neural circuits involving the anterior cingulate cortex and supplementary motor area (Bonini et al., 2014; Dehaene, Posner, & Tucker, 1994; Gentsch, Ullsperger, & Ullsperger, 2009). Although frequently used as a measure of individual and group differences related to psychopathology (Moser, Moran, Schroder, Donnellan, & Yeung, 2013; Olvet & Hajcak, 2008; Seer, Lange, Georgiev, Jahanshahi, & Kopp, 2016), the ERN can be confounded by task interpretation and task performance, with larger amplitudes when focusing on accuracy (Gehring, Goss, Coles, Meyer, & Donchin, 1993) and in individuals who make fewer errors (Fischer, Klein, & Ullsperger, 2017). Differences in the ERN across development (Tamnes, Walhovd, Torstveit, Sells, & Fjell, 2013) and aging (Beste, Dziobek, Hielscher, Willemssen, & Falkenstein, 2009a; Nieuwenhuis et al., 2002) can be difficult to interpret because different age groups, such as young children, need different tasks to maintain similar levels of engagement and difficulty in order to minimize confounds related to differences in motivation and performance accuracy (Lewis & Stieben, 2004). And while many theories implicate the ERN in adaptive behavior (Holroyd & Coles, 2002b; Ullsperger, Danielmeier, & Jocham, 2014), incremental trial-by trial adaptation is not robustly testable or readily observable in tasks with categorical response options. Although response latencies demonstrate post-error slowing that are often considered to be adaptive, post-error slowing may instead reflect orienting responses to infrequent events (Notebaert et al., 2009). The ERN appears to reflect the activity of a generic error detection system that responds similarly to errors committed by the hands, feet (Holroyd, Dien, & Coles, 1998), or eyes (Van’t Ent & Apkarian, 1999), to continuous motor errors that are independent of decision-making (Gallea, Graaf, Pailhous, & Bonnard, 2008; Maurer, Maurer, & Muller, 2015), and even after errors that are not committed by the individual, if the individual is responsible for correcting the error (Gentsch et al., 2009). Given this flexibility in the type of errors detected, it may be possible to leverage more controllable tasks to evoke an ERN-like potential.

A sudden disturbance to standing balance evokes a frontocentral negativity called the balance N1 that resembles the ERN in several ways (Payne et al., 2019b), but it is unknown whether these potentials share variance across individuals. The balance N1 is frontocentrally distributed and localized to the supplementary motor area (Marlin, Mochizuki, Staines, & McIlroy, 2014; Mierau, Hulsdunker, & Struder, 2015), with simultaneous activation of the anterior cingulate cortex (Peterson & Ferris, 2018, 2019). Both the balance N1 and the ERN increase in amplitude with the perceived consequence of an error, with a larger balance N1 evoked at the edge of a raised platform compared to ground level (Adkin, Campbell, Chua, & Carpenter, 2008; Sibley, Mochizuki, Frank, & McIlroy, 2010), and a larger ERN when errors are more costly or being evaluated and judged (Hajcak, Moser, Yeung, & Simons, 2005a; Kim, Iwaki, Uno, & Fujita, 2005). Both increase with the extent of the error: the balance N1 increases with larger disturbances (Mochizuki, Boe, Marlin, & McIlRoy, 2010; Payne, Hajcak, & Ting, 2019a; Staines, McIlroy, & Brooke, 2001), and the ERN increases with the extent of an error, such as when an error is committed with both the wrong finger and the wrong hand (Bernstein, Scheffers, & Coles, 1995). Both the ERN and balance N1 depend on attention, decreasing in amplitude when distracted by a secondary task (Klawohn, Endrass, Preuss, Riesel, & Kathmann, 2016; Little & Woollacott, 2015; Quant, Adkin, Staines, Maki, & McIlroy, 2004b). Both the ERN and balance N1 decrease in amplitude when errors are more expected, such as when a countdown precedes a familiar balance disturbance (Adkin, Quant, Maki, & McIlroy, 2006; Mochizuki, Sibley, Cheung, & McIlroy, 2009b), or when errors occur more frequently with increased task difficulty (Van der Borght, Houtman, Burle, & Notebaert, 2016). Both ERPs register in the theta frequency range in time-frequency analyses (Luu, Tucker, & Makeig, 2004; Peterson & Ferris, 2018, 2019; Varghese et al., 2014). Like the ERN, the balance N1 can be evoked in a variety of tasks, such as shoving the torso (Adkin et al., 2006); sudden release of support (Mochizuki et al., 2010); sudden tilt of the floor while standing (Ackermann, Diener, & Dichgans, 1986); or sudden slip-like movements of the floor while standing (Payne et al., 2019a), walking (Dietz, Quintern, & Berger, 1985b), or sitting (Mochizuki, Sibley, Cheung, Camilleri, & McIlroy, 2009a; Staines et al., 2001). The balance N1 appears to be consistent with a generic system for error detection as it does not differ in timing or amplitude between conditions in which either the arms or the legs are used to recover balance (Mochizuki et al., 2009a). While the balance N1 and the ERN share many features and factors, it is unknown whether they share variance across individuals, such that individuals with a larger balance N1 also have a larger ERN.

If the balance N1 were to reflect the individual differences in error processing that are indexed by the ERN, there are several advantages to a balance paradigm. Balance errors are intrinsically motivating and evoke an involuntary balance-correcting behavioral reaction (Jacobs & Horak, 2007a) that does not require instruction on how to perform or perceive the task, making it possible to avoid potential bias due to task instruction or interpretation. Further, balance errors are experimentally controllable, allowing the exact same series of errors to be repeated across individuals (Adkin et al., 2006; Welch & Ting, 2014), thereby eliminating the need for participants and groups to spontaneously commit comparable sequences of mistakes. Balance-recovery behavior also demonstrates rapid context-dependent trial-by-trial adaptation of behavior at multiple time scales (Quintern, Berger, & Dietz, 1985) that is readily observable across a range of outcome variables such as the extent of muscle activation and body movement (Horak & Nashner, 1986; Welch & Ting, 2014).The balance N1 can be evoked during standing balance in nearly anyone able to stand without an assistive device, from toddlers (Berger, Horstmann, & Dietz, 1990; Berger, Quintern, & Dietz, 1987) through the elderly (Duckrow, Abu-Hasaballah, Whipple, & Wolfson, 1999; Payne, McKay, & Ting, 2022; Payne, Palmer, McKay, & Ting, 2021). Further, the balance N1 is quite large, being robustly observable on individual trials (Ditz, Schwarz, & Muller-Putz, 2020; Payne et al., 2019a), and therefore has the potential to yield better psychometric properties with fewer trials than the ERN.

As a preliminary assessment of whether the balance N1 and the ERN share variance across individuals, we added measurement of the ERN into two separate, ongoing investigations of the balance N1. These two studies included small groups of young or older adults who were given different protocols of slip-like balance disturbances in which the floor would suddenly move as if a rug were being pulled out from beneath the standing participant. Both studies included the exact same arrow flanker task, which was completed twice at the end of the lab visit. The two iterations of the flanker task differed in response modality, with responses entered by either the hand or the feet. Within each group, we compared the ERNs between hand and foot response modalities to establish consistency of the ERN across two objectively similar tasks. We evaluated internal consistency of all ERPs, and finally, compared the balance N1 to the ERN from each version of the flanker task to assess whether the balance N1 and the ERN share variance across individuals.

## 2. Methods

### 2.1 Study 1 Methods

### 2.1.1 Participants

Twenty-one healthy young adults (age 25±5 years, range 19-38, 12 female) were recruited from the community surrounding Emory University. The protocol was approved by the Institutional Review Board of Emory University, and all participants were informed of the study procedures and provided written consent before participation. Different analyses of the balance N1 potential in this population are previously published (Payne & Ting, 2020a; Payne & Ting, 2020c).

### 2.1.2. Balance Task

Participants were exposed to a series of 48 translational support-surface balance perturbations that were unpredictable in timing and magnitude (**FIGURE 1**). The support-surface, (i.e., the floor, and thus the feet) moved backward in all perturbations, resulting in a relative forward lean of the body. Three perturbation magnitudes were used to ensure unpredictability of the perturbation characteristics. The small perturbation was identical across participants (7.7 cm, 16.0 cm/s, 0.23 g). The medium (12.6-15.0 cm, 26.6-31.5 cm/s, 0.38-0.45 g) and large (18.4-21.9 cm, 38.7-42.3 cm/s, 0.54-0.64 g) perturbations were scaled by participant height to control for the effect of participant height when using support-surface translations to evoke the balance N1 (Payne et al., 2019a) and to ensure that the more difficult perturbations were mechanically similar across body sizes. To prevent fatigue, 5-minute rest breaks were enforced when the full duration of the perturbation series was expected to exceed 16 minutes, with additional breaks allowed upon request. Not counting these breaks, the duration of the perturbation series was 17±2 minutes across participants. Inter-trial-intervals between perturbation onsets were 22±3 s, with perturbations manually triggered when the EEG baseline was relatively quiescent, approximately 5-15 s after the participant returned to a stable, upright posture.

**Figure 1.**
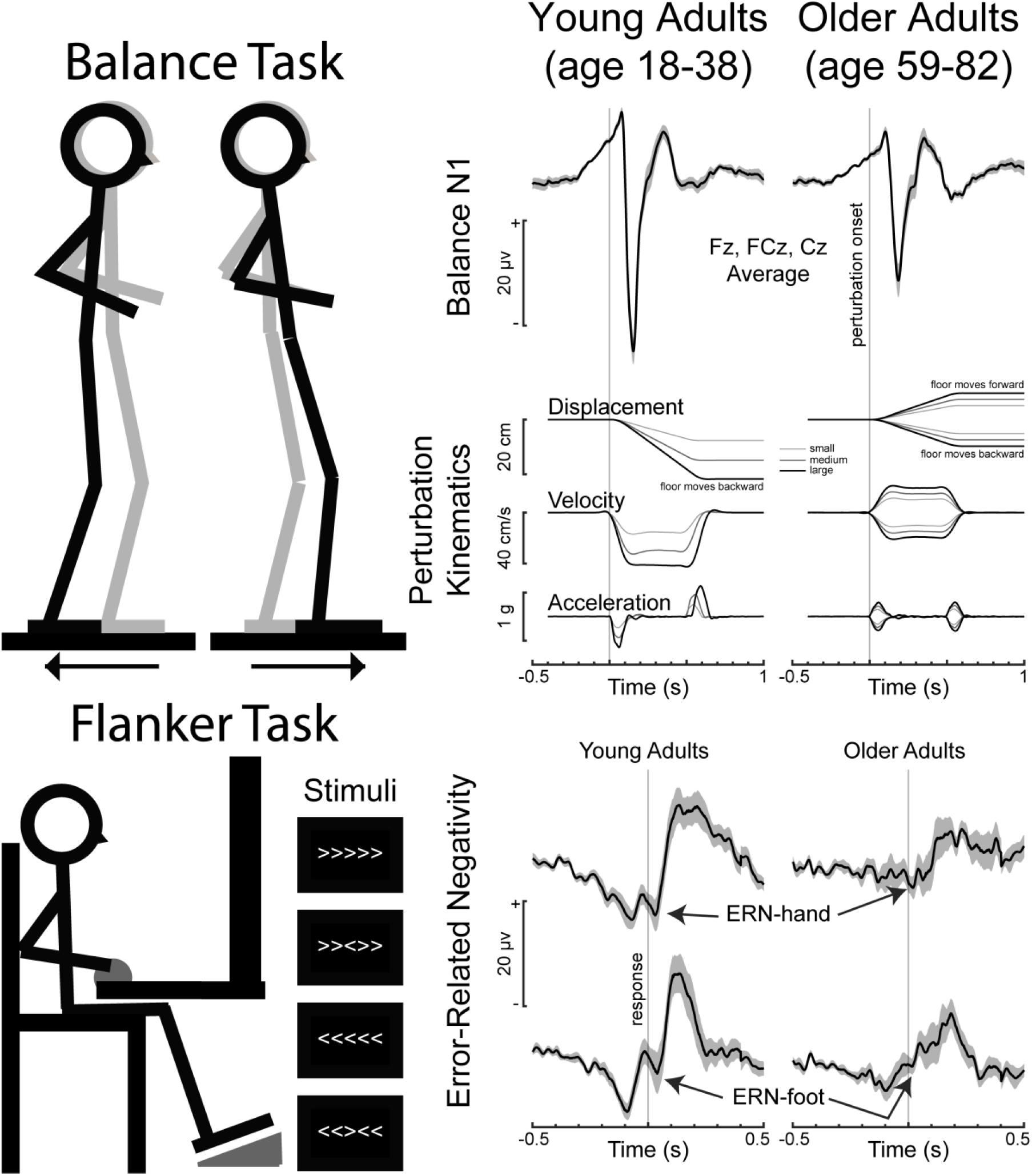
Experimental Tasks. (Top panel) A schematic depicts the balance task along with grand-averaged balance N1s and perturbation kinematics from young and older adult populations. (Bottom panel) A schematic depicts the flanker task, with the two response entry devices depicted in gray. Grand-averaged error-related negativities (ERNs) are shown for the hand and foot response modalities of the flanker task in both populations. ERP data in all panels is plotted positive-up and averaged across Fz, FCz, and Cz, with the standard error of the mean across participants shaded. Note that the young adults received only backward perturbations while the older adults received both backward and forward perturbations of smaller magnitudes. Axes are matched to enable comparisons between young and older adults, as well as between balance and flanker ERPs.

Participants were harnessed to the ceiling for safety using a harness that provided no weight support during perturbations. The harness could suddenly catch a participant if they began to fall toward the ground, but this did not occur in either of the populations reported here. Participants were instructed to cross their arms in front of their chest and focus their gaze at a picture of a mountain landscape 4.5 meters ahead during the initial platform motion. On half of trials, participants were asked to recover balance by taking a single step forward. On the remaining trials participants were asked to recover balance without taking a step, but participants occasionally stepped involuntarily in response to the larger perturbations despite efforts to keep their feet in place. Stepping instructions were varied randomly between blocks of 6 trials, with each block containing two replicates of each perturbation magnitude in random order. Although we previously reported small differences in the balance N1 across perturbation magnitudes in some individuals (Payne et al., 2019a; Payne & Ting, 2020a), and small changes in the balance N1 with stepping at the largest perturbation magnitude (Payne & Ting, 2020c), the present analyses will collapse across all trial types to maximize and balance the number of trials included in the measurement of the balance N1. Due to failure to save the EEG data, data from the perturbation series is unavailable for one participant.

### 2.1.3 Flanker Tasks

After the perturbation series and a 5-minute rest break, participants performed two versions of an arrowhead flanker task (Eriksen & Eriksen, 1974) in counterbalanced order using Presentation software (Neurobehavioral Systems, Inc., Albany, California). The two versions differed in response modality. In one version, participants responded to stimuli by clicking the left or right mouse button using the pointer and middle fingers of the hand of their choice. In the other version, participants responded to stimuli by releasing a foot pedal under the ball of the left or right foot, with the pedals otherwise remained pressed throughout the task. The tasks were otherwise identical with one exception that a message to “Please ensure both foot pedals are depressed” was inserted between stimuli in the foot response version if the foot pedals were not engaged at the time the next stimulus was supposed to be displayed.

In both task versions, participants were shown five arrowheads in each trial, and were instructed to respond as quickly and accurately as possible according to the direction of the central arrowhead (**FIGURE 1**). Stimuli were balanced between compatible (“>>>>>“ and “<<<<<“) and incompatible (“>><>>“ and “<<><<“) conditions in random order. Each stimulus was displayed for 200 ms, and the interval between offset of one stimulus and onset of the subsequent stimulus varied randomly between 2300-2800 ms, unless delayed by failure to engage foot pedals as described above. In such a case, the message would disappear once the pedals were pressed, and the next stimulus was displayed 2300-2800 ms later. Participants first completed a supervised practice block of 11 trials, which could be repeated if participants still did not understand the task. After the practice block, each task consisted of up to 10 blocks of 30 trials (up to 311 total trials), with each block initiated by the participant. The task was set to terminate early if 21 errors were obtained (Meyer et al., 2013), as further increases in reliability are small beyond 20 errors (Fischer et al., 2017). This termination condition was met more often than not (15/19 cases for the hand response modality and 16/18 cases for the foot response modality), as any accuracy below 93% will reach 21 errors within 300 trials. In attempt to maintain accuracy between 75-90%, messages were displayed between blocks stating, “Please try to be more accurate,” “Please try to respond faster,” or “You’re doing a great job,” according to the accuracy of the preceding block.

Three participants completed only the hand response version of the task and two participants completed only the foot response version of the task due to changes in the experimental protocol across the first five participants. The remaining sixteen participants completed both versions of the flanker task counterbalanced in order.

### 2.1.4. EEG Collection

EEG data were collected during all three tasks using a 32-channel active electrode system (ActiCAP, Brain Products, Germany) placed according to the international 10-20 system. Electrodes TP9 and TP10 were removed from the cap and placed directly on the skin over the left and right mastoid bones for offline re-referencing. Electrodes were prepared with conductive electrode gel (SuperVisc HighViscosity Electrolyte-Gel, Brain Products) using a syringe that gently abraded the scalp to improve impedances. Impedances at Cz and mastoid electrodes were generally below 10 kOhm before the start of data collection. Vertical EOG was collected to correct for blink and eye movement artifacts using bipolar passive electrodes (E220x, Brain Products), placed above and below the right eye with a forehead reference. EOG electrodes were prepared with high-chloride abrasive gel (ABRALYT HiCl, High-chloride-10% abrasive electrolyte gel, Brain Products). EEG and EOG data were amplified on an ActiCHamp amplifier (Brain Products) and sampled at 1000 Hz following a 24-bit A/D converter and 20 kHz online anti-aliasing low-pass filter. The EEG system also recorded data from a 3D accelerometer placed on the head to monitor for the possibility of motion artifacts during the balance task.

### 2.1.5. EEG Pre-Processing

EEG data were high-pass filtered at 1 Hz with forward and backward passes of third-order Butterworth filters to ensure zero lag, mean-subtracted within channels, and then similarly low-pass filtered at 25 Hz. EEG data were re-referenced to the average of the mastoids and epoched around perturbation onset for the balance perturbations, and around response entry for the flanker task (detailed below) at Fz, FCz, Cz, and the vertical EOG channel. The Gratton and Coles algorithm (Gratton, Coles, & Donchin, 1983) was applied to correct for blink and eye movement artifacts at Fz, FCz, and Cz using the vertical EOG channel. Due to limited numbers of trials within any given trial type (e.g., errors, nonstepping responses to large backward perturbations, etc.), trial types were not distinguished in the correction of eye artifacts, but data from the three tasks were processed separately.

### 2.1.6. Balance Task ERPs

Filtered and re-referenced EEG data from the perturbation series were epoched in the period of 400 ms before perturbation onset, defined as the onset of perturbation acceleration (**FIGURE 1**), until 2000 ms after perturbation onset. After eye artifact correction (described above), subject-averaged ERPs were created by averaging the EEG data across all trials within each participant, separately for Fz, FCz, and Cz. No perturbation trials were excluded from analysis for any reason as the signal-to-noise ratio of the balance N1 is larger than most typical artifacts, and a quiet baseline was manually confirmed prior to perturbation onset. Additionally, due to the nature of the perturbations applied at the feet, significant head motion did not occur until after the balance N1 (**FIGURE 2**), and thus head acceleration data were not assessed further. The balance N1 was quantified as the most negative amplitude in subject-averaged ERPs between 100-250 ms after perturbation onset relative to the mean of a baseline period of 50-150 ms before perturbation onset.

**Figure 2.**
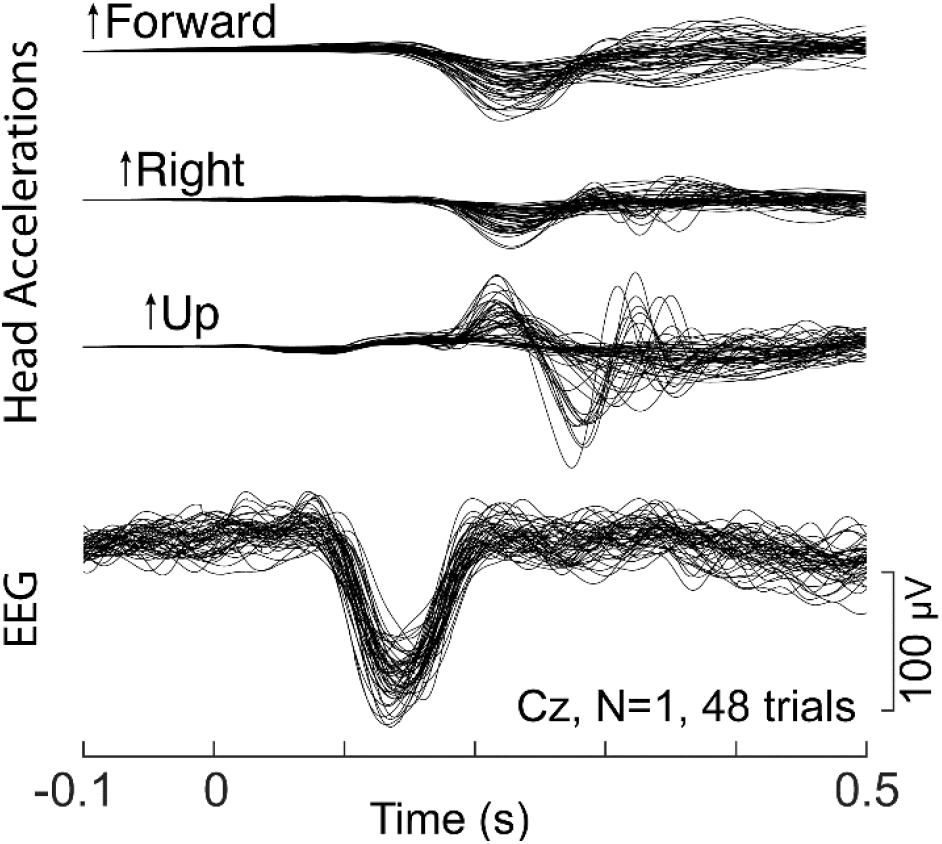
The balance N1 is not a motion artifact. 3D head accelerations and balance N1s are shown for all trials in the balance task from one example participant in the young adult group. Note that head accelerations to the left in this example arise from anticipatory postural adjustments preceding a step to recover balance with the right foot.

### 2.1.7 Flanker Task ERPs

EEG data were processed separately for hand and foot response versions of the flanker task. Filtered and re-referenced EEG data were epoched in 1000 ms segments centered on response entry at Fz, FCz, and Cz. Trials were discarded from analyses according to the following automated rejection criteria: <0.5 μV difference for over 100 ms, >50 μV difference between consecutive samples, >300 μV difference across the epoch, reaction times more than 3 standard deviations from the participant’s average reaction time, within-trial range in voltage more than 2 standard deviations above the participant’s average, or within trial variance across samples more than 2 standard deviations above the subject’s mean. Subject-averaged ERPs were created using the remaining error trials, separately at Fz, FCz, and Cz. To mimic the scoring approach of the N1, the ERN was measured in the ERP from error trials as the most negative point between 50 ms before response to 100 ms after response relative to the average of a baseline of 300-500 ms prior to response entry. Similar results were obtained when ERPs were measured as the mean of a 50 ms window centered around the peak. For simplicity, ERNs from the two different response modalities will be referred to as the ERN-hand and the ERN-foot.

All of the young adults had at least six artifact-free error trials, and thus no young adults were excluded for an insufficient number of error trials. Average numbers of error trials entered into analyses for each electrode and task are reported in the results.

### 2.1.5. Statistical Analyses

Internal consistency was assessed in terms of the split-half reliability. Split-half reliability was assessed by comparing the amplitudes of the ERPs created by separately averaging the even and odd trials that went into the subject averages. Specifically, the split-half reliability was taken as the Pearson’s correlation coefficient between the amplitudes measured from the even and odd averaged waveforms across participants, which was subsequently corrected using the Spearman-Brown prophecy formula (Cassidy, Robertson, & O’Connell, 2012; Warrens, 2017).

Pearson product-moment correlations were then used to test for associations between the ERN-hand and ERN-foot, as well as between the balance N1 and each of the ERNs. Variables deviating from a normal distribution (Shapiro-Wilk p<0.05) were transformed to a normal distribution using boxcox.m in MATLAB (MathWorks Inc.) prior to calculation of statistics. Scatter plots display original, untransformed data values along with corrected statistics as appropriate.

A principal component analysis (PCA) was used to measure the variance shared across the three ERPs. Specifically, PCA (pca.m in MATLAB) was applied to the three ERP amplitudes (separately for each electrode). The variables were scaled to unit variance before PCA to balance the contributions of each ERP. Missing data were handled using the alternating least squares algorithm (’ALS’ option in pca.m), enabling inclusion of all participants, even those who did not complete all three tasks. Because this algorithm does not result in a deterministic solution, we report the result across ten iterations of PCA. As all three ERPs loaded strongly onto the largest principal component, the proportion of the total variance accounted for by that component was used as a measure of the variance shared across the three ERPs.

Additionally, to assess whether there was variance unique to the ERNs that was not shared with the balance N1, we performed an additional Pearson product-moment correlation between the residual variance in the ERN-hand and ERN-foot after using regression and subtraction to remove the component of variance that each shared with the balance N1 (Meyer, Lerner, De Los Reyes, Laird, & Hajcak, 2017). Measuring the correlation between these residual variances measures the variance that is shared between the two ERNs beyond what they share balance N1.

### 2.2 Study 2 Methods

### 2.2.1. Participants

Twenty healthy older adults (age 70±7 years, range 59-82, 7 female) were recruited from the community surrounding Emory University. The protocol was approved by the Institutional Review Board of Emory University, and all participants were informed of the study procedures and provided written consent before participation.

Unless specified below, all methods and procedures were identical to Study 1. The flanker tasks were identical across studies. The early termination condition of 21 errors was met for 7/19 cases for the hand response modality and 13/19 cases for the foot response modality. One older adult was unable to perform either version of the flanker task due to inability to perceive the brief 200 ms stimuli. Another older adult was excluded from analyses for near chance accuracy (53% accuracy in hand version, 47% accuracy in foot version), and an additional two older adults were excluded from analyses of the ERN-hand due to fewer than 6 artifact-free error trials. Additionally, due to poor signal quality at Cz (resulting from repeated electrode pop-off during the experimental session), data at the Cz electrode was replaced with the average of electrodes C1 and C2 in one individual across all three tasks.

### 2.2.2. Balance task

Participants were exposed to a series of 48 translational support-surface balance perturbations that were unpredictable in timing, magnitude, and direction. This differs from Study 1, in which the exclusively backward direction of perturbations was predictable.

Perturbations were balanced between forward and backward perturbation directions, and between three perturbation magnitudes in block-randomized orders (**Figure 1**). The small perturbation was identical across participants (5.1 cm, 11.1 cm/s, 0.15 g), while the medium (7.0-7.4 cm, 15.2-16.1 cm/s, 0.21-0.22 g) and large (8.9-9.8 cm, 19.1-21.0 cm/s, 0.26-0.29 g) perturbations were scaled according to participant height. These perturbation magnitudes were smaller than those used in Study 1, and unlike Study 1, participants were asked to try not to take a step in all trials. A 5-minute break was enforced half-way through the perturbation series, with additional breaks allowed upon request. Not counting breaks, the perturbation series lasted 18±2 minutes, and inter-trial intervals between perturbation onsets were 23±10 s.

### 2.2.3. Statistical analyses

All analyses were performed as described in Study 1. In addition to the analyses described in Study 1, we also used one-tailed *t*-tests to assess whether ERNs were smaller in the older adults compared to the younger adults to establish consistency with existing studies (Beste, Willemssen, Saft, & Falkenstein, 2009b; Nieuwenhuis et al., 2002). While the balance N1 is also smaller in older adults (Duckrow et al., 1999), a similar comparison in the present data would be confounded by the smaller perturbation magnitudes in the older group. Although the same balance perturbations could have been used between the young and older adult groups, these studies were originally designed to address different experimental questions related to the balance N1 potential (Payne et al., 2022; Payne et al., 2021; Payne & Ting, 2020a; Payne & Ting, 2020c).

## 3. Results

### 3.1. Study 1 Results

In the hand version of the flanker task, young adults (N=19) committed an average of 19±3 errors, with 90±4% accuracy, and reaction times of 385±23 ms (errors 337±35 ms, correct 391±24 ms). After trial rejections, 17±3 error trials were included in the measurement of the ERN-hand (Fz 16±3, FCz 17±3, Cz 17±3). ERN-hand amplitudes were -9.3±5.5, -9.1±3.8, and -7.0±3.2 μV at Fz, FCz, and Cz. In the foot version of the flanker task, young adults (N=18) committed an average of 21±2 errors, with 89±4% accuracy, and reaction times of 405±35 ms (errors 331±26 ms, correct 414±36 ms). After trial rejections, 17±2 error trials were included in the measurement of the ERN-foot (Fz 16±3, FCz 17±2, Cz 18±2). ERN-foot amplitudes were -7.5±5.7, -7.4±5.3, and -6.3±4.6 μV at Fz, FCz, and Cz.

ERNs showed good reliability and were associated in amplitude across the hand and foot flanker tasks. Split-half reliabilities at Fz, FCz, and Cz were 0.90, 0.86, and 0.79 for the ERN-hand, and 0.86, 0.84, and 0.80 for the ERN-foot (**FIGURE 3A**). The ERN-hand and ERN-foot were associated in amplitude across the young adults (**FIGURE 3B**, N=16, Fz p=0.0001 r=0.817, FCz p=0.055 r=0.487, Cz p=0.33 r=0.259).

**Figure 3.**
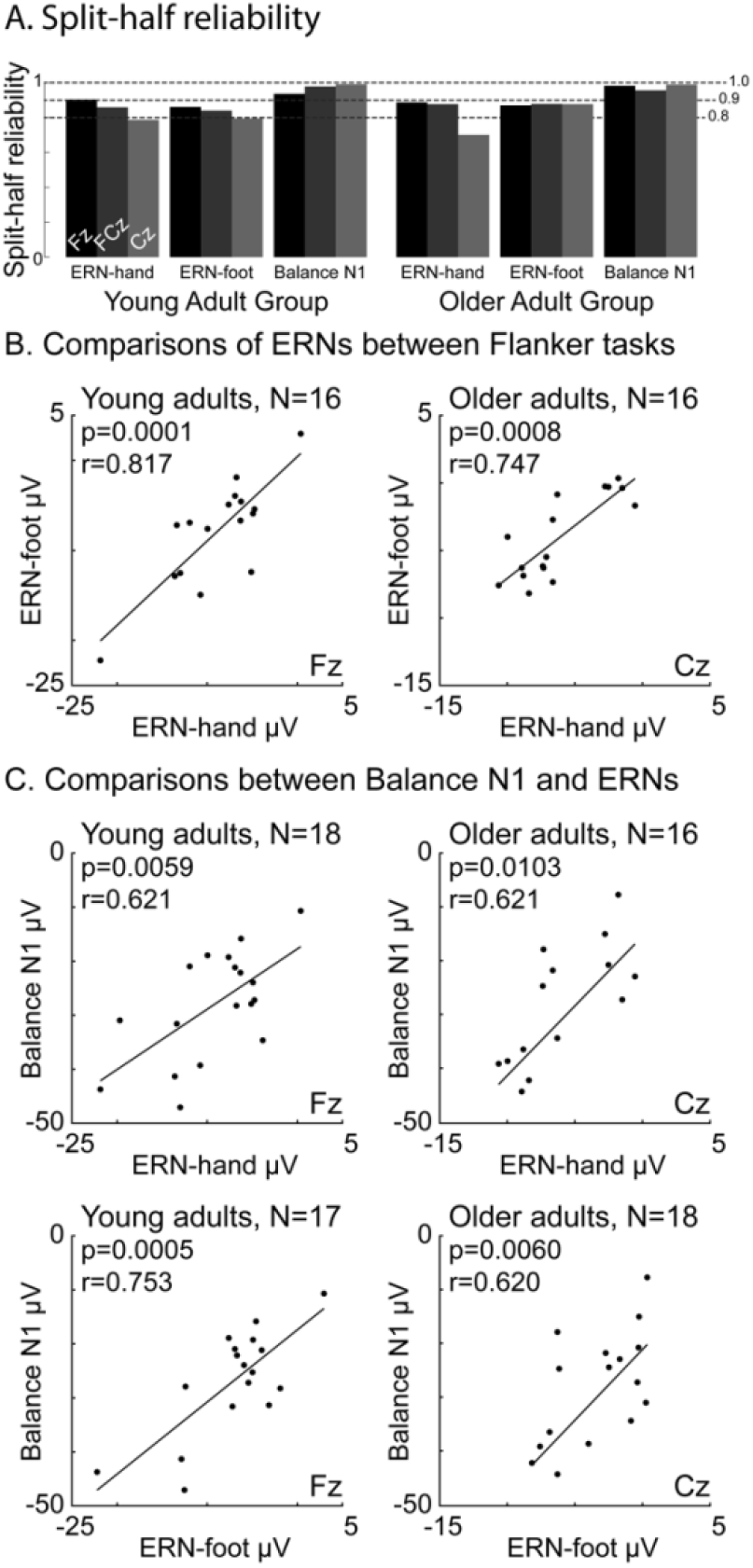
Comparison of ERPs across tasks. (A) Bar plots show split-half reliability of all ERPs. (B) Scatter plots show associations in the error-related negativity (ERN) between the hand and foot response versions of the flanker task. (C) Scatter plots show associations between the balance N1 and the ERNs from the hand and foot response versions of the flanker task. Note that this figure depicts results at different electrodes for each group because the groups differed in the site of strongest associations, but results are reported for three frontocentral electrodes for both groups.

The balance N1 showed excellent reliability and was associated in amplitude with the ERNs. Split-half reliabilities at Fz, FCz, and Cz for the balance N1 were 0.94, 0.98, and 0.99 (**FIGURE 3A**). Balance N1 amplitudes in the young adults (N=19) were -28.1±9.6, -43.8±12.5, and -54.1±18.0 at Fz, FCz, and Cz. Balance N1 amplitudes were associated with the ERN-hand (**FIGURE 3C**, N=18, Fz p=0.0059 r=0.621, FCz p=0.0317 r=0.507, Cz p=0.062 r=0.449) and the ERN-foot (N=17, Fz p=0.00048 r=0.753, FCz p=0.0204 r=0.556, Cz p=0.224 r=0.331).

Variance was shared across the three evoked potentials with additional variance specific to the flanker ERNs in the young adults. All three ERPs strongly loaded onto the largest principal component, which accounted for the majority of individual differences in ERP amplitudes (**FIGURE 4A** Fz 75±10% of variance, FCz 69±11%, Cz 62±5%). When the correlation to the balance N1 was regressed out of the ERN-hand and the ERN-foot, the residual variances maintained an association at Fz (**FIGURE 4B** Fz p=0.020 r=0.593, FCz p=0.072 r=0.478, Cz p=0.327 r=0.272).

**Figure 4.**
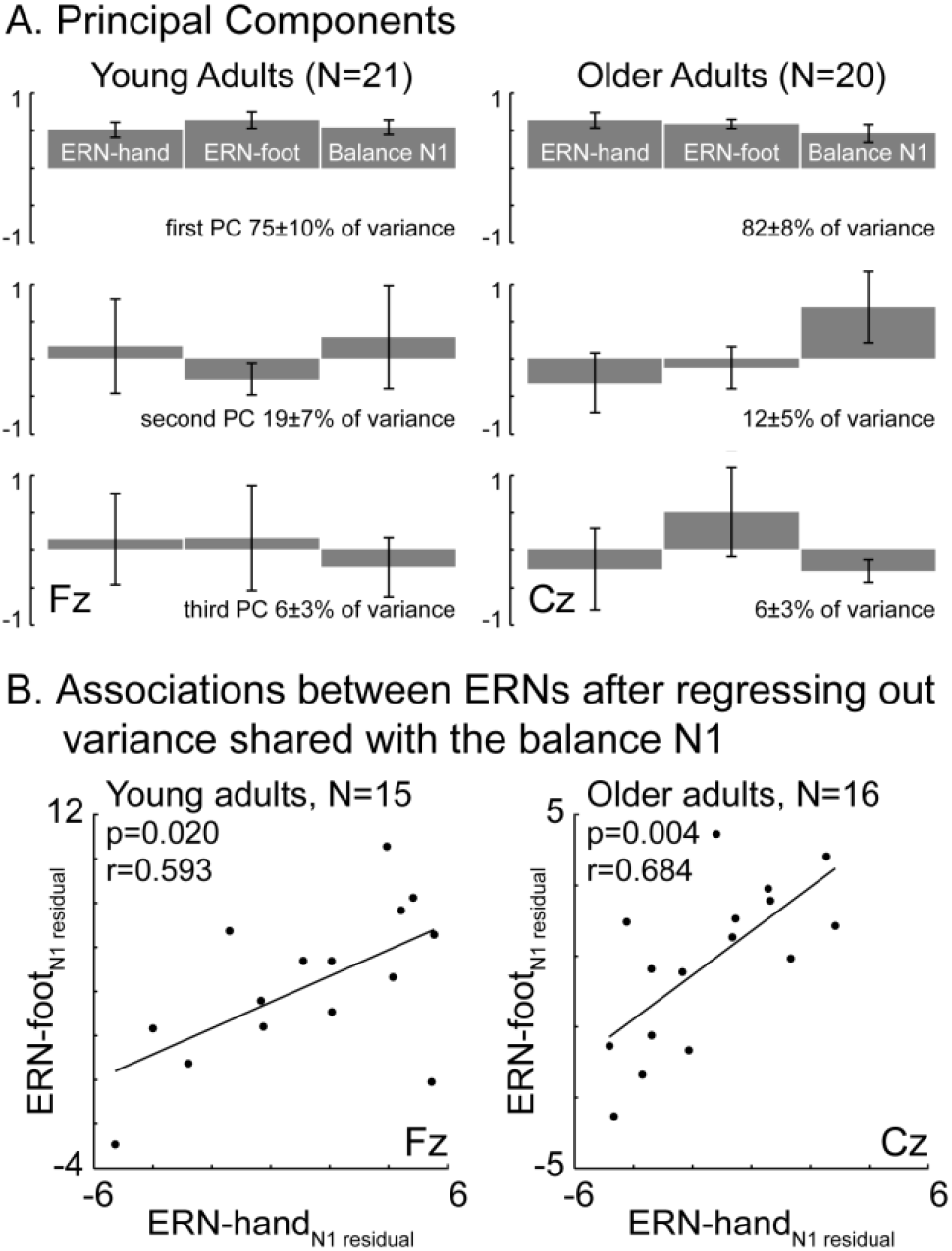
Variance shared across ERPs. (A) The principal components are shown for young adults at Fz (left) and for older adults at Cz (right). (B) Scatter plots show the association that remains between the ERN-foot and ERN-hand after the variance each variable shared with the balance N1 has been regressed out (for the young adults at Fz on the left and the older adults at Cz on the right).

### 3.2 Study 2 Results

In the hand version of the flanker task, older adults (N=16) committed an average of 16±5 errors, with 93±4% accuracy, and reaction times of 503±149 ms (errors 459±298 ms, correct 509±144 ms). After trial rejections, 14±5 error trials were included in the measurement of the ERN-hand (Fz 13±5, FCz 14±5, Cz 14±5). ERN-hand amplitudes were -5.3±4.5, -6.1±3.0, and -6.1±3.2 μV at Fz, FCz, and Cz. In the foot version of the flanker task, older adults (N=18) committed an average of 21±1 errors, with 85±7% accuracy, and reaction times of 462±58 ms (errors 359±51 ms, correct 482±60 ms). After trial rejections, 18±2 error trials were included in the measurement of the ERN-foot (Fz 17±3, FCz 18±2, Cz 18±2). ERN-foot amplitudes were -4.7±4.6, -4.0±3.3, and -3.7±3.1 μV at Fz, FCz, and Cz.

ERNs showed good reliability at most sites and were associated in amplitude across the hand and foot flanker tasks. Split-half reliabilities at Fz, FCz, and Cz were 0.89, 0.88, and 0.70 for the ERN-hand, and 0.87, 0.88, and 0.88 for the ERN-foot (**FIGURE 3A**). The ERN-hand and ERN-foot were associated in amplitude across the older adults (**FIGURE 3B**, N=16, Fz p=0.243 r=0.309, FCz p=0.011 r=0.615, Cz p=0.00081 r=0.747).

The balance N1 showed excellent reliability and was associated in amplitude with the ERNs. Split-half reliabilities at Fz, FCz, and Cz for the balance N1 were 0.98, 0.96, and 0.99 (**FIGURE 3A**). Balance N1 amplitudes in the older adults (N=20) were -22.6±11.5, -29.7±13.0, and -32.9±14 at Fz, FCz, and Cz. Balance N1 amplitudes were associated with the ERN-hand (**FIGURE 3C** N=16, Fz p=0.704 r=0.103, FCz p=0.092 r=0.435, Cz p=0.0103 r=0.621) and the ERN-foot (N=18, Fz p=0.109 r=0.391, FCz p=0.00491 r=0.737, Cz p=0.0060 r=0.620).

Variance was shared across the three evoked potentials, with additional variance specific to the flanker ERNs in the older adults. All three ERPs strongly loaded onto the largest principal component, which accounted for the majority of individual differences in ERP amplitudes (**FIGURE 4A** Fz 59±8% of variance, FCz 68±10%, Cz 82±8%). When the correlation to the balance N1 was regressed out of the ERN-hand and the ERN-foot, the residual variances maintained an association at Cz (**FIGURE 4B** Fz p=0.273 r=0.292, FCz p=0.087 r=0.441, Cz p=0.004 r=0.684).

The older adult group had smaller ERNs than the younger group in both the hand (Fz p=0.032, FCz p=0.008, Cz p=0.21) and foot (Fz p=0.057, FCz p=0.014, Cz p=0.027) versions of the flanker task.

## 4. Discussion

We previously suggested that the balance N1 and the ERN may share underlying mechanisms based on a number of shared features and factors (Payne et al., 2019b); based on results from the current study, we now add that they share variance across individuals in two small samples that vary in age across the adult lifespan. Internal consistency reliabilities were good for the ERNs (0.70-0.90 across groups and electrodes) and excellent for the balance N1 (0.94-0.99). We found that ERNs evoked in an arrow flanker task are strongly correlated across hand and foot response modalities, and that these ERNs both have moderate to strong correlations with the balance N1. In both groups, a single principal component strongly loaded on all three ERPs, suggesting that the majority of individual differences are shared across these ERPs. However, there remains a significant component of variance shared between the ERN-hand and ERN-foot beyond what they share with the balance N1. It is unclear whether this component of variance is task-specific (e.g., variance that would not be shared with the ERN evoked by a go/no-go task), or something fundamentally related to error processing that is not evoked by a sudden balance disturbance. Important next steps will be to assess both the ERN and the balance N1 in more rigorous experimental designs with larger sample sizes, and to determine whether the variance shared between the balance N1 and the ERN relate to individual differences of interest to research in development and psychopathology. If the balance N1 were to reflect error processing mechanisms indexed by the ERN, balance paradigms offer several advantages in terms of experimental control over errors.

To our knowledge, this is the first study directly examining the correlation in amplitude between the ERN elicited across hand and foot response modalities. Internal consistency measures for the ERN were above 0.80 at most sites for both groups, indicating good reliability, consistent with previous reports (Fischer et al., 2017; Riesel et al., 2013). Prior work has shown that both hand and foot errors elicit an ERN that does not differ in localization between hand and foot response modalities in young adults (Holroyd et al., 1998). We now demonstrate that ERN amplitudes are strongly correlated between these response modalities across the adult lifespan. Hand and foot ERNs were more correlated at frontal electrode sites in the young adults and more correlated at central sites in the older adults—although the reason for this is unclear. The ERN-foot was frontally maximal in both younger and older adults, whereas the ERN-hand in the older adults was largest at central sites. Although unexpected, this is not unprecedented, as one study demonstrated that the scalp distribution of ERN amplitude associations differs across pairings between Go/NoGo, Stroop, and flanker tasks (Riesel et al., 2013). Additionally, our data replicate prior work demonstrating that the ERN amplitude is smaller in older compared to younger adults (Beste et al., 2009b; Nieuwenhuis et al., 2002), but comparisons across the present groups may be confounded by differences in motivation, accuracy, and fatigue or demand characteristics at the end of different experimental paradigms.

We believe this is the first study to assess psychometric properties of the balance N1, as well as the first study to test whether the balance N1 shares variance in amplitude with the ERN across individuals. The higher internal consistencies for the balance N1 compared to the ERNs may relate to both a greater amplitude of the balance N1 and a greater number of trials available to assess the balance N1. While the correlations and principal component analyses demonstrate that the majority of variance was shared across the three ERPs, the residualized regression demonstrates that there remains a significant portion of variance shared between the flanker task ERNs beyond what they share with the balance N1. Further investigations will need to assess whether the balance N1 reflects the group and individual differences of interest to research in development (Tamnes et al., 2013) and psychopathology (Moser et al., 2013; Olvet & Hajcak, 2008; Seer et al., 2016). Future studies could examine, for instance, whether anxious individuals have a larger balance N1. One notable difference is that the balance N1 is localized to the supplementary motor area (Marlin et al., 2014; Mierau et al., 2015) while the ERN is localized to the nearby anterior cingulate cortex, although both of these areas are active during both ERPs (Bonini et al., 2014; Peterson & Ferris, 2018, 2019). While this may be a meaningful difference between the two ERPs, it is possible that the nature of the stimulus determines the relative recruitment of these reciprocally connected cortical areas, similar to how the stimulus content determines the relative recruitment of the reciprocally connected cognitive and affective divisions within the anterior cingulate cortex (Bush, Luu, & Posner, 2000), which may bias localization under the assumption of a single source.

There are important limitations to consider. While replicating a large effect size of correlation across two independent samples adds support to the suggestion that a balance disturbance can evoke an ERN-like potential, it will be necessary to confirm these findings in larger samples, allowing other variables like age and sex to be considered. The ERN was not the primary focus of these studies (Payne et al., 2022; Payne et al., 2021; Payne & Ting, 2020a; Payne & Ting, 2020c), and was always collected at the end of the session, after the physically active balance task. Although acute exercise does not necessarily influence the ERN (Themanson & Hillman, 2006), it is possible that the fixed task order had some effect on the current results. Using the same perturbation series across populations would have enabled more comparisons across groups, but the older adults received much easier perturbations because they were recruited as a control population to be compared against Parkinson’s disease in another study (Payne et al., 2022). Although the excellent internal consistency reliability of the balance N1 suggests that a reliable estimate could be obtained using fewer trials, the possibility for habituation across initial trials should always be carefully considered, especially where differences in habituation could present an additional source of individual differences (Payne et al., 2019a). While the even-odd method of splitting the data would balance any potential effect of initial habituation, the split-half reliability of the balance N1 was similar when splitting instead between the first and second half of trials, but this may not be the case when a non-randomized perturbation series is used (Quintern et al., 1985). Finally, although perturbations applied at the feet do not move the head during the balance N1, we strongly recommend the use of active electrodes and measurement of head acceleration, especially in more accessible methods of perturbing the trunk (Adkin et al., 2006; Mochizuki et al., 2010), which may accelerate the head at shorter latencies.

If the balance N1 were to reflect the same error processing mechanisms indexed by the ERN, then specific advantages of balance paradigms could be leveraged to investigate error processing. Bias due to instruction and interpretation can be avoided in balance paradigms because balance recovery behavior begins with a stereotyped involuntary behavior requiring no prior instruction (Jacobs & Horak, 2007a). There is no reliance on participants to commit comparable sequences of mistakes, as the exact same series of balance errors can be delivered to every individual (Adkin et al., 2006; Welch & Ting, 2014). Additionally, the balance N1 is sufficiently large to observe on individual trials (**FIGURE 2**) and may offer better psychometric properties than the ERN evoked in speeded-response tasks (**FIGURE 3**).

Although slip-like disturbances in which the floor suddenly moves require specialized equipment, other methods of disturbing balance can be implemented much more easily. Balance disturbances that rely on gravity, applied as a sudden release of support, can be as straightforward as manually removing a pin (Mochizuki et al., 2010), and allow for more precise time-locking than systems relying on motors (Payne et al., 2019a). If the balance N1 and the ERN arise from overlapping neural mechanisms, their relationship might also provide insight into comorbidities between balance and anxiety disorders (Balaban, 2002; Balaban & Thayer, 2001; Bolmont, Gangloff, Vouriot, & Perrin, 2002; Jacob et al., 1997; Yardley & Redfern, 2001) and treatment, such as balance training that alleviates anxiety in young children (Bart et al., 2009) and the benefit of psychotherapy to balance disorders (Schmid, Henningsen, Dieterich, Sattel, & Lahmann, 2011; Yardley & Redfern, 2001).

## Author notes

## Acknowledgements

Study data were collected and managed using a Research Electronic Data Capture (REDCap) database hosted at Emory University (Harris et al., 2019; Harris et al., 2009).

## CRediT statement

LHT provided funding; AMP, LHT, and GH were involved in the conceptualization, methodology, and interpretation of the findings; AMP collected and analyzed the data, drafted the manuscript, and prepared figures; LHT and GH provided feedback throughout data analyses and revision of the manuscript and figures.

